# Prediction of synonymous corrections by the BE-FF computational tool expands the targeting scope of base editing

**DOI:** 10.1101/2020.01.06.890244

**Authors:** Rabinowitz Roy, Abadi Shiran, Almog Shiri, Offen Daniel

## Abstract

Base editing is a genome-editing approach that employs the CRISPR/Cas system to precisely install point mutations within the genome. A cytidine or adenosine deaminase enzyme is fused to a deactivated Cas and converts C to T or A to G, respectively. The diversified repertoire of base editors, varied in their Cas and deaminase proteins, provides a wide range of functionality. However, existing base-editors can only induce transition substitutions in a specified region determined by the base editor, thus, they are incompatible for many point mutations. Here, we present BE-FF (Base Editors Functional Finder), a novel computational tool that identifies suitable base editors to correct the translated sequence erred by a given single nucleotide variation. Even if a perfect correction of the single nucleotide variation is not possible, BE-FF detects synonymous corrections to produce the reference protein. To assess the potential of BE-FF, we analysed a database of human pathogenic point mutations and found suitable base editors for 60.9% of the transition mutations. Importantly, 19.4% of them were made possible only by synonymous corrections. Moreover, we detected 298 cases in which pathogenic mutations caused by transversions were potentially repairable by base editing via synonymous corrections, although it had been thought impractical. The BE-FF tool and the database are available at https://www.danioffenlab.com/be-ff.

**GRAPHICAL ABSTRACT:** 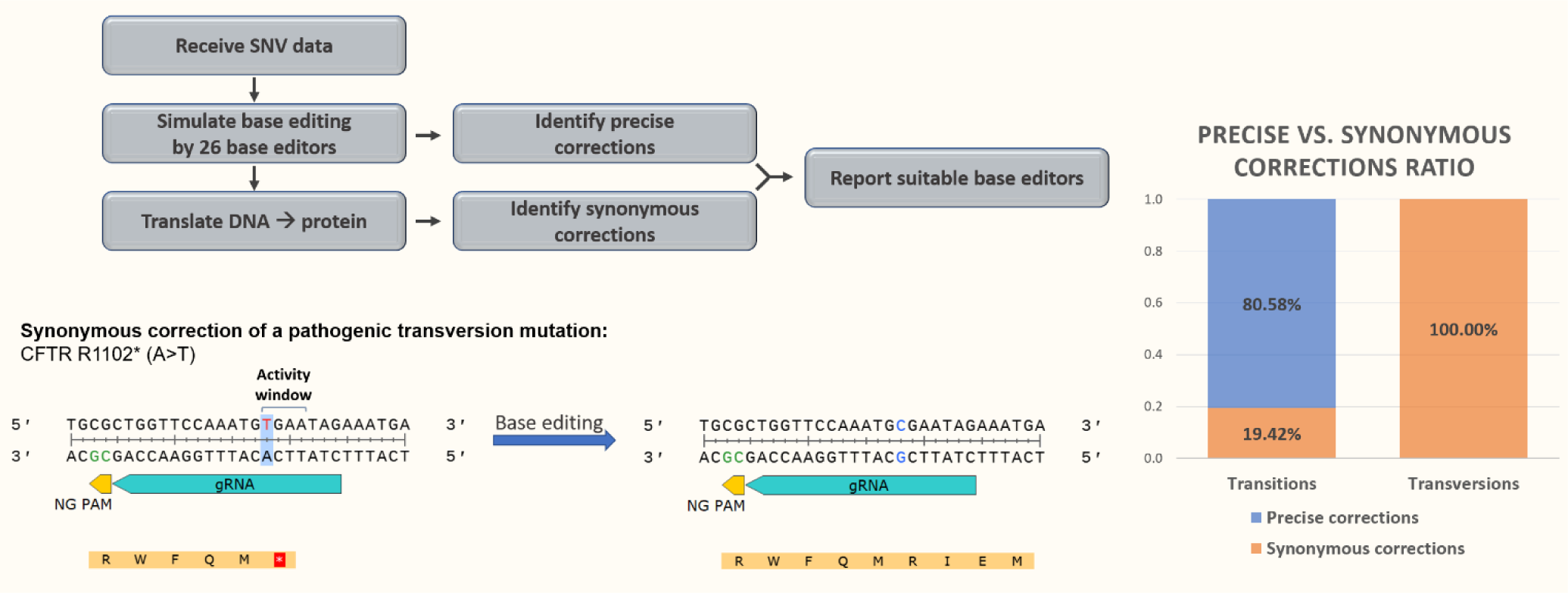

## INTRODUCTION

Base editors (BEs) allow programmable genome editing in terms of a single nucleotide transition; purine to purine and pyrimidine to pyrimidine (A↔G and C↔T, respectively)(1, 2). The base editing technology employs the clustered regulatory interspaced short palindromic repeats (CRISPR)/Cas system to deliver a deaminase protein to precise genomic loci, as directed by the guide-RNA (gRNA)(3, 4). The first BE (BE1) was introduced by Komor et al. (1). This BE utilizes a cytidine deaminase enzyme fused to a catalytically deactivated Cas (dCas)(1), a Cas protein that contains mutations within its RuvC and HNH endonuclease domains (D10A and H840A) leading to the inability of the Cas protein to perform DNA cleavage. While the dCas protein lacks its endonuclease ability, it retains the competence to navigate through the genomic DNA to the designated locus(5). Many more variants have been devised since then (table 1) and can be categorized to two main types: cytosine BEs (CBEs) which convert C to T and adenine BEs (ABEs) that convert A to G. The conversion by CBEs occurs via deamination of cytidine and the generation of uridine that acts as thymidine in base pairing(1). ABEs utilize an adenosine deaminase enzyme to perform adenosine deamination, resulting in an inosine. During translation inosine acts as guanosine(6), hence the activity of ABE yields an A to G transition(2). By targeting the complementary strand, it is possible to indirectly convert G to A by CBE and T to C by ABE. Taken together, CBEs and ABEs are capable of performing all the converting combinations of transition substitutions. While point mutations account for 58% of disease-causing genetic variants in humans, transition substitutions comprise 61% of the pathogenic point mutations(7).

**Table 1:**
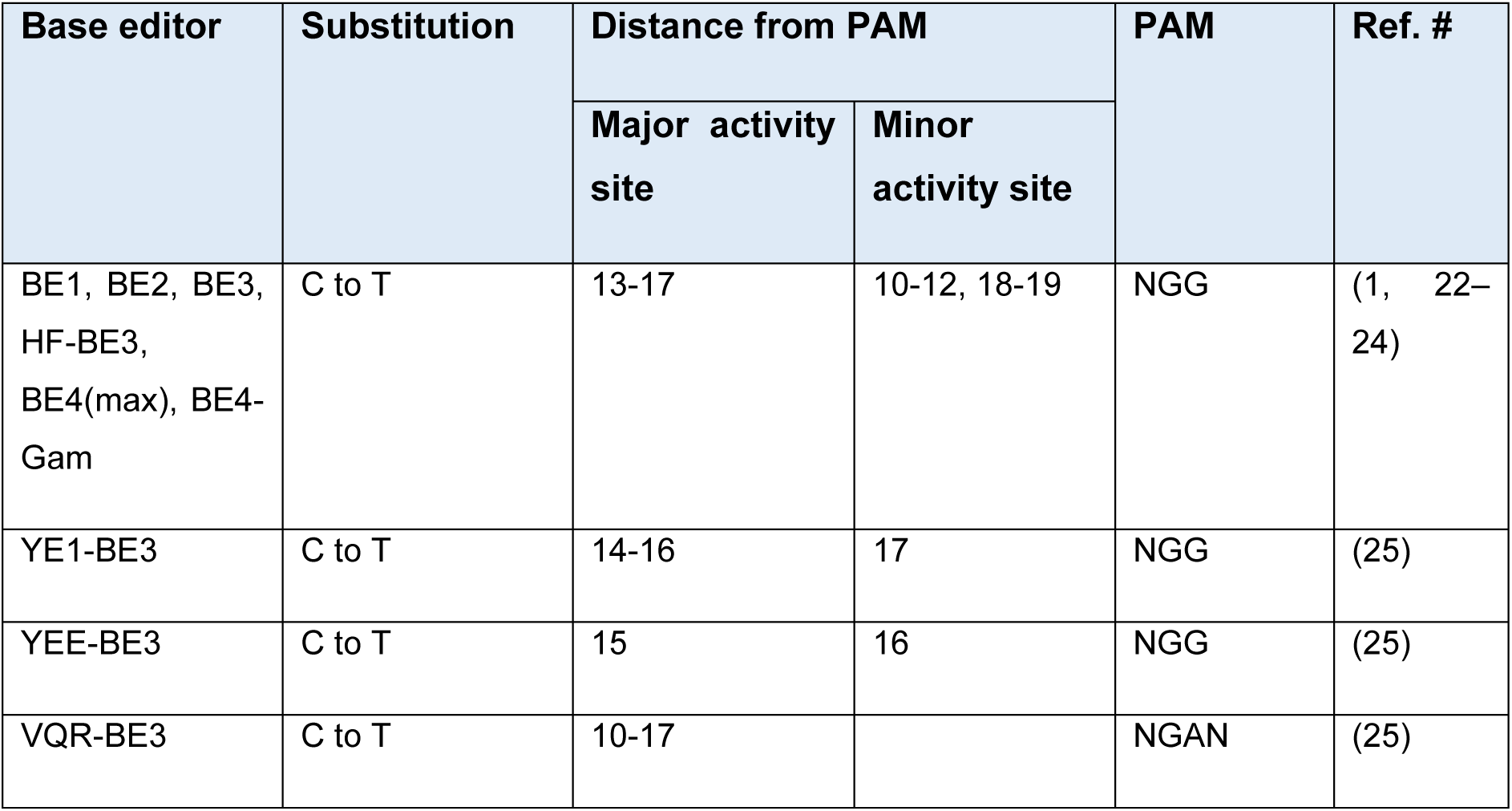
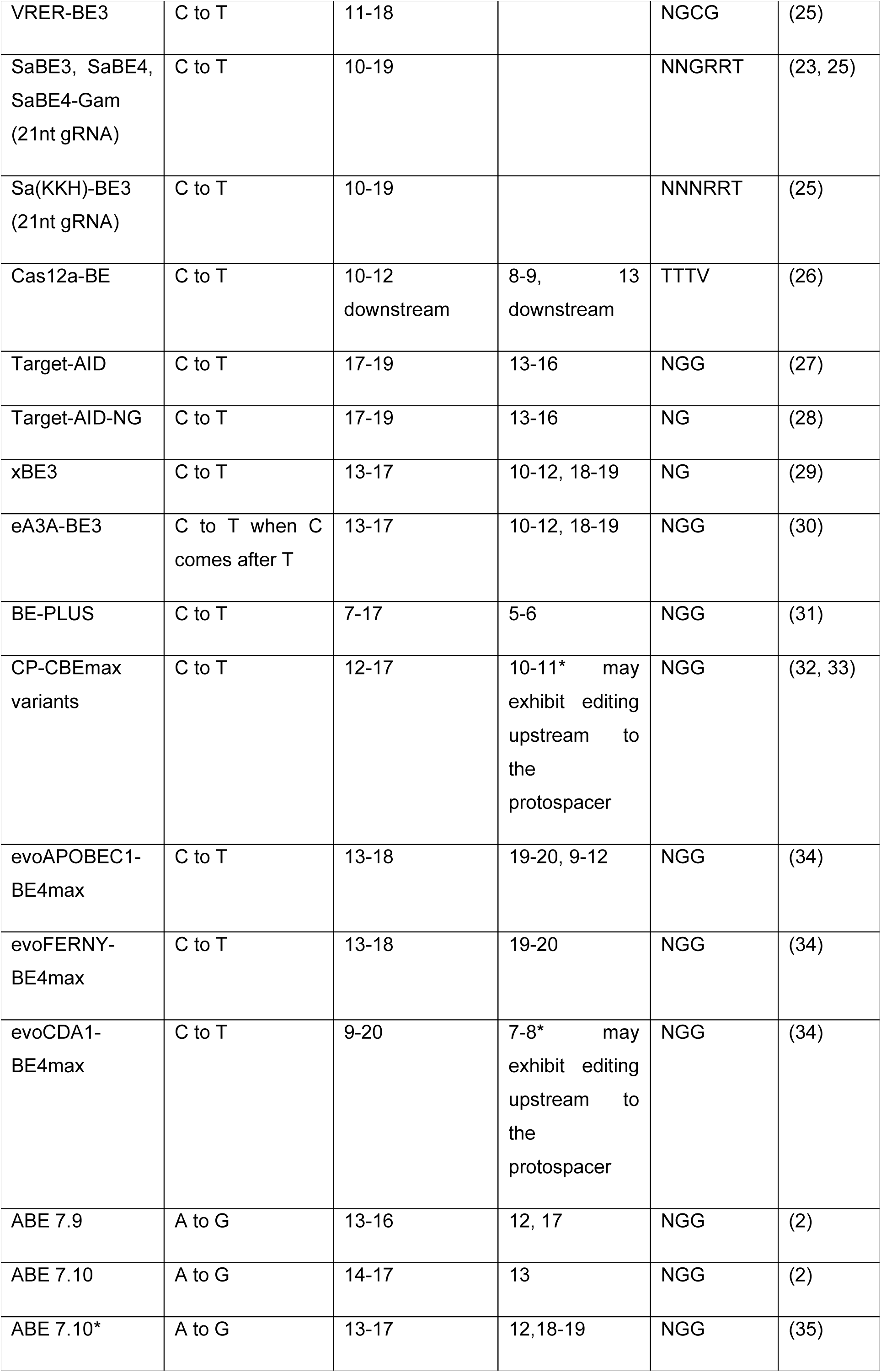
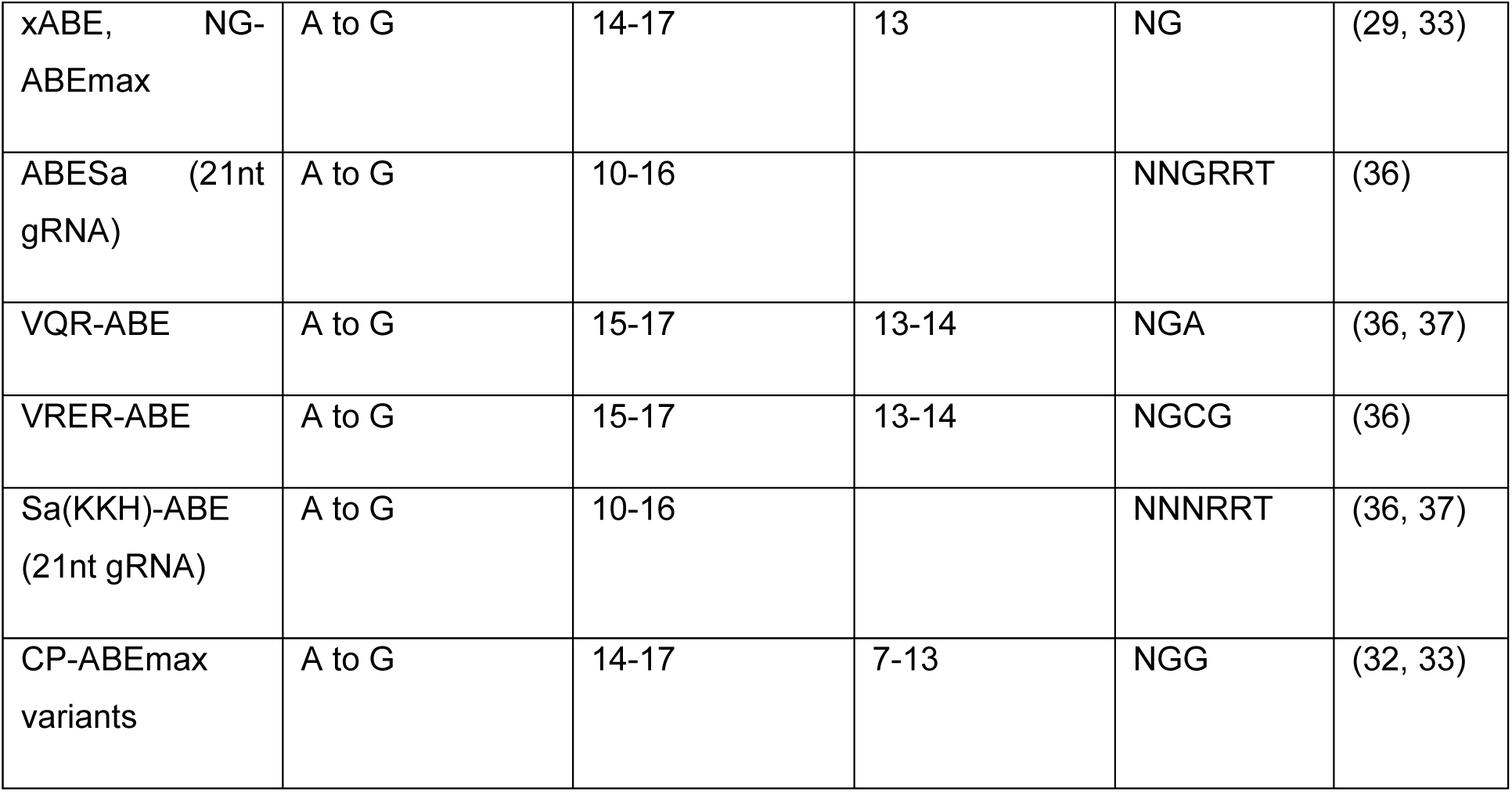
BEs repository.

Notably, contrary to other CRISPR mediated gene editing methods, base editing does not involve DNA double-strand breaks (DSBs); thus, conferring a higher degree of safety as DSBs consequences include error-prone mutagenic repair pathways (alternative end joining and single-strand annealing)(8), p53 activation(9, 10), large deletions and rearrangements(11), integration of foreign genomes at the target site(12) and more. Furthermore, compared to the homology directed repair (HDR) pathway that is thought to be the only precise resolution amongst DSB repair pathways, base editing is both more efficient(1) and allows editing of post-mitotic cells that are unable to undergo DSB-mediated HDR(13, 14). A diverse toolbox of CBEs and ABEs is essential for developing treatments based on base editing for disease-associated point mutations. As natural Cas proteins are being discovered and synthetic variants are constantly being reported, so does the base editors toolbox expands with novel CBEs and ABEs. A pivotal consideration in gRNA design in general and base editing in particular, is the protospacer adjacent motif (PAM) limitation. The PAM is a short sequence within the target DNA that has an essential role in DNA binding of the Cas protein. The motif must be flanking the target sequence as directed by the gRNA, downstream or upstream according to the Cas type (type II and type V, respectively) in order to induce DNA cleavage by the Cas protein(15). As the PAM determines the binding site of the Cas protein to the DNA, it dictates the activity window region of the BE. Therefore, targeting a particular nucleotide is narrowed by the presence of a PAM in a precise distance from the activity window as determined by the BE. Each BE has a major activity window, where base editing occurs most efficiently, and minor activity window(s) in which the BE exhibits some degree of editing in significantly lower rates. All the target nucleotides (C or A) within the major activity window will undergo base editing. Consequently, if a target nucleotide is flanked by the same nucleotide, both will be edited and an unintended mutation may be introduced to the DNA (bystander base editing), instead of correction of the gene. In some cases, bystander base editing leads to a synonymous mutation compared to the intended sequence and may be accepted as a successful base editing outcome (e.g., ACTCTA [Thr,Leu] to ATTTTA [Ile, Leu] where threonine is the variant and isoleucine is the reference amino acid). A BE is comprised of the 3 basic elements of Cas and deaminase proteins and a linker fusing the 2 proteins. Thus, each BE has its own unique features: PAM compatibility, gRNA length, orientation relative to the PAM, affinity to the target sequence, target nucleotide (C or A), efficiency, activity window width and its distance from PAM, off-targets, protein size and more.

Since single-nucleotide variants (SNVs) naturally vary in genetic context, a diverse range of BEs is essential to precisely adjust to a given SNV. In base editing experimental design, one should consider the properties of the available BEs alongside their basic fit to perform transition of the target nucleotide. Due to the large selection of BEs (table 1) and the complexity of identifying proper BEs to a target site, the necessity of a computational tool arises. gRNA design and off-targets prediction tools are available for general purposes such as gene-knockouts(16–19). However, current tools use reference genomes as a template, while point mutations and patient-derived cells differ from the reference genome and therefore such tools are not suitable for designing base editing experiments for treating point mutations. Moreover, such tools are not customized for base editing and thus, do not take under consideration BEs’ activity window and amino acids (AA) sequence. Existing tools that are base editing oriented, do not match suitable BEs for specific SNVs (20), or lack the possibility to examine the translation outcome of the edited sequence(21). To magnify the potential of base editing in treating as many cases as possible, the utilization of multiple Cas varieties and the ability to translate DNA sequences and compare the editing outcome are needed. To that end, we developed BE-FF, a tool that receives SNVs data, analyzes the reference and variant sequences and their translated outcomes and matches the suitable BEs out of 26 unique BEs. To assess the potential of base editing as a therapeutic approach for genetic diseases, we demonstrate the efficiency of BE-FF on a dataset of human pathogenic and likely-pathogenic SNVs. Furthermore, we established the BE-FF DB, a comprehensive database that includes pathogenic SNVs that can be edited via BE.

## RESULTS

### A database of human pathogenic point mutations and their applicable BEs

First, we sought to assess the potential of base editing to treat human pathogenic point mutations. To that end, we assembled a large collection of pathogenic SNVs and identified four possible scenarios of successful base editing: a) Precise correction in which the resulting edited DNA sequence resembles the desired reference sequence (figure 1a). b) Multiple bases synonymous correction of the target nucleotide together with synonymous editing of bystander nucleotides, meaning that the resulting protein sequence resembles the reference protein sequence although the DNA sequences differs (figure 1b). c) On-target synonymous correction is the case in which the target nucleotide is converted into a different nucleotide other than the reference one, though this conversion rescues the protein sequence (figure 1c). d) Base editing of a bystander nucleotide within the codon of the pathogenic point mutation (bystander synonymous correction) rescues the protein sequence without editing the pathogenic SNV. This case is mostly advantageous for transversion point mutations. While BEs are unable to reverse the DNA sequence of transversion mutations to match the reference sequence, editing a bystander nucleotide results in a proper AA substitution that matches the reference protein sequence (figure 1d). While the first two scenarios allow the correction of transition mutations, the last two scenarios also allow the correction of transversion mutations as they exploit codon degeneracy of several AAs.

**Figure 1:**
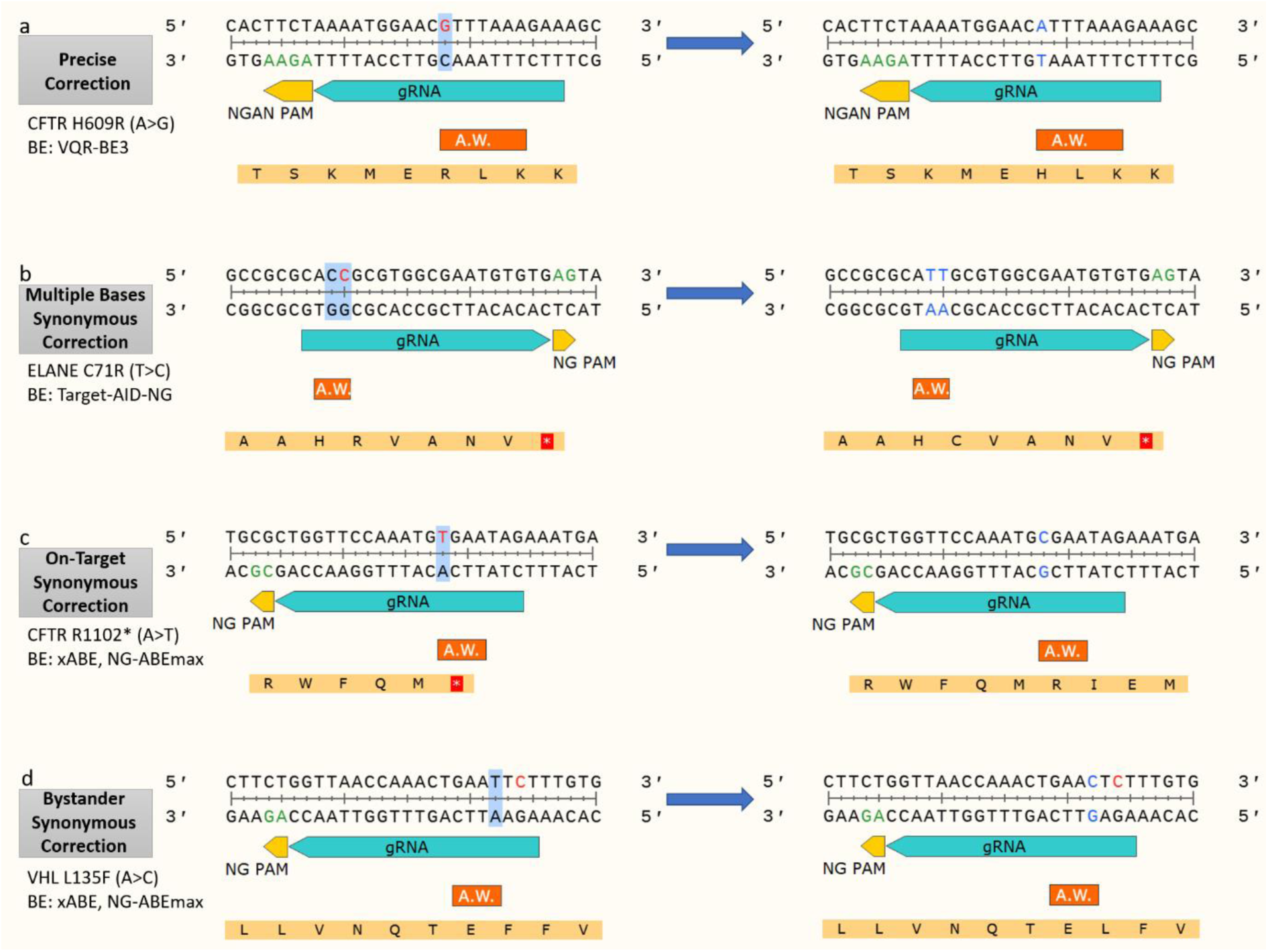
Base editing correction scenarios. The gRNA and PAM sequences appear in bright blue and yellow, respectively. The major activity window of the base editor is shown as A.W. in orange. The left sequences represent the pathogenic SNV (red) sequences and the right sequences represent the simulated base-edited (blue) sequences. The target nucleotides within the activity window are marked with blue background. a) Precise correction: the CFTR H609R transition mutation perfectly repaired by VQR-BE3. b-d) Synonymous correction scenarios. The resulting DNA sequence does not match the reference allele; however, the translated protein sequence matches. b) Multiple bases synonymous correction: in addition to the target nucleotide, a bystander nucleotide lies within the activity window and undergoes base editing. c) On-target synonymous correction: the variant nucleotide is not restored to the reference nucleotide. The resulted codon is encoded to the reference AA. d) Bystander synonymous correction: the target nucleotide remains intact while a bystander editing restores the protein sequence.

SNPs data was obtained from NCBI’s dbSNP(38) (SNPs with either pathogenic or likely-pathogenic clinical significance). Insertion, deletion and multiple base substitution mutations were filtered out due to their lack of compatibility for base editing. The SNPs dataset on which we performed our analysis contained 43,504 SNPs; 27,098 transitions and 16,406 transversions (supplementary file 1). In theory, any transition mutation could be corrected given that it is positioned within the major activity window of a suitable BE. Indeed, we found that 60.9% of the transitions can be repairable (figure 2a-b), while 19.4% of them could be corrected only by synonymous corrections (supplementary figure s1). Even though no existing BE can repair a transversion SNV, we detected 298 pathogenic SNPs caused by transversion (1.8% of the total transversion mutations) for which the resulting AA could be reversed, i.e., corrected by inducing a transition editing in the SNP site or a bystander site to generate a synonymous codon (figure 2b).

**Figure 2:**
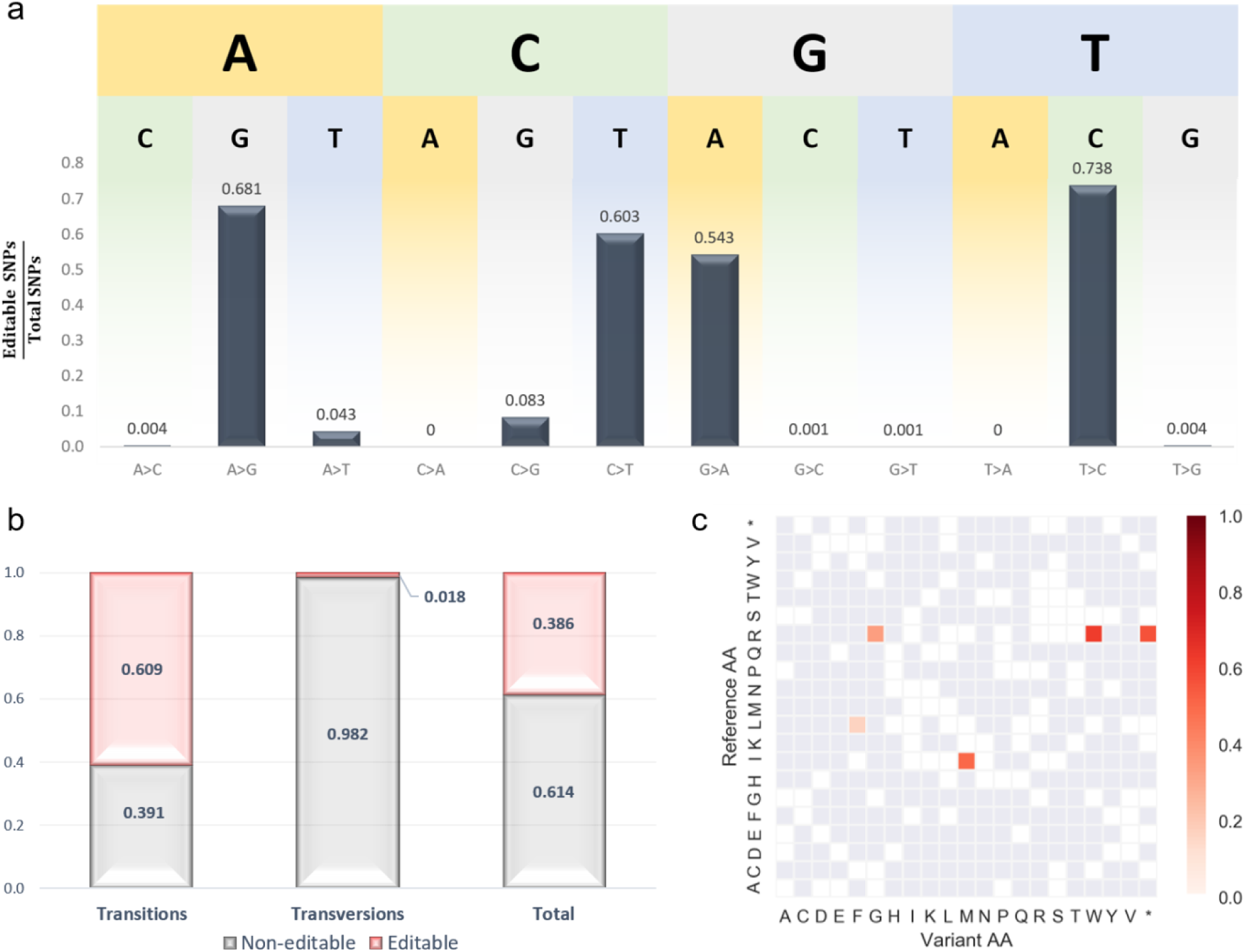
Human pathogenic mutations repaired by base editing. a) Ratios of editable SNVs out of total SNVs for all substitution combinations. b) Ratios of editable and non-editable SNVs for transition mutations (60.9% and 39.1%, respectively) and transversion mutations (1.8% and 98.2%, respectively). In total, 38.6% of the tested pathogenic SNVs were repairable. c) Heatmap representation of AA substitutions caused by transversions repaired by base editing (y axis - reference AAs, x axis – variant AAs). Five AA substitutions were repairable: I>M, L>F, R>G, R>W and R>* (52%, 16%, 33%, 63% and 57% of the total of the transversion-derived mutations for each, respectively).

We further sought to assess the frequencies at which AA substitutions that were caused by transversion are reversible. Our results reveal that the following AA substitutions, caused by transversions, could be reversed via base editing: I>M, L>F, R>G, R>W and R>* (52%, 16%, 33%, 63% and 57% of the total of the transversion-derived mutations for each, respectively; figure 2c). As expected, the majority of possible synonymous correction of transversions are those of AAs that are encoded by 6 codons (leucine and arginine). The third position of the isoleucine codons is considered the solely threefold degenerate site, meaning that a substitution in that position to 2 out of 3 alternative nucleotides results in no change of the AA. Only a change of H (A/C/U) to G would result in an I to M missense mutation. Therefore, such I to M mutations caused by transversions (T to G or C to G) could be reversed by CBEs (G to A). Serine, although encoded by 6 codons as well, could not be recovered by base editing due to the difference between its two sets of codons that do not allow the flexibility of leucine and arginine (UCN and AGY; where N complies to A/C/G/U thus generating a synonymous mutation in the flexible site a-priori and Y complies to C/U thereby only a transversion substitution could have corrected this site).

### BE-FF: Base Editors Functional Finder – a web tool that identifies BEs to correct SNVs

We established a web tool that receives SNV data and matches suitable BEs to correct the variation. The web tool receives data as a user manual input, fetches data by rsID or an uploaded batch mode file. Together with the flanking regions of the SNV in the DNA sequence, the reading frame of the sequence is utilized to translate the sequence. All 26 BEs (table 1) are available and examined to match the query. The tool does not limit the repertoire of BEs according to the base substitution. Thus, for any given SNV, an attempt to match any of the BEs is made to detect ones that perform precise correction as well as synonymous correction. The reverse-complement sequences are also considered for correction of the coding sequence. While a precise correction requires a full match of both the DNA and AA sequences, synonymous corrections are considered positive when only the AA sequences match and the DNA sequences do not. It is assumed that precise corrections are favorable in most cases over synonymous corrections and therefore the output divides the results into 2 parts, precise corrections and synonymous corrections. BE-FF supports user defined BE properties to allow researchers the utilization of novel or unpublished BEs.

### Comparison to available tools

We compared BE-FF to two available base-editing design tools in several parameters and report the differences in a comparative table (table 2).

**Table 2:** Base editing tools comparative table.

|  | **BE-FF** | **BE-Designer**(20) | **BEable-GPS**(21) |
| --- | --- | --- | --- |
| **Multiple BE varieties** | 17 CBEs and 9 ABEs | Limited (3 CBEs, 1 ABE) | Limited (only CBEs) |
| **Customized user defined BE support** | V | V (limited to predefined PAM list) | V (limited to CBEs only) |
| **Translate editing outcome and detect synonymous corrections** | V | V | X |
| **Support transversion synonymous corrections** | V | V | X |
| **Support editing of the complement sequence for correction of the coding sequence** | V | V | V |
| **Simple user interface** | V | V | V |
| **Off-targets assessment** | X | V | X |
| **Design BE gRNA for specific SNVs** | V | X | V |
| **Multiple BEs comparison for a given SNV** | V | X | V |
| **Targeting approach** | Detects suitable BEs to reverse a given SNV | Shows the predicted base editing outcome for a given sequence by a user defined BE | Target region can be specified. Shows the base editing outcome of selected BEs |
| **Input format** | Fetch by SNP ID,<br>uploaded file of<br>multiple SNVs or<br>user manual input | User manual input<br>or uploaded file of<br>multiple sequences | User manual<br>input |

## DISCUSSION

In this study we report the feasibility of editing several transversion-mediated AA substitutions by base editing, although mutations caused by transversions are thought to be excluded from the scope of base editing. Our data demonstrate that in most cases, the following substitutions: I>M, L>F, R>G, R>W and R>*, are repairable via base editing and encourage considering base editing as an ideal approach for correcting such mutations. Amongst the repairable transversion mutations are SNVs associated with varied conditions including cystic fibrosis, deafness, Fanconi anemia and more, emphasizing the significance of BE-FF in base editing gRNA design. We utilize the properties of 26 base editors (17 CBEs and 9 ABEs) that vary in their Cas proteins, deaminase enzymes and linkers and therefore provide a broad toolbox to perform base editing. The development of novel BEs contributes to the expansion of the base editing toolbox and advances base editing towards future therapeutics and research applications. We compared BE-FF to existing tools and find its utility valuable due to its compatibility with both CBEs and ABEs, the ability of translation and comparing the translated sequences to identify synonymous corrections, vast repertoire of BEs and the simple user interface. BE-FF is ideal for finding base editing solutions to repair specific point mutations. A recent study by Anzalone et al. reports a novel method termed Prime editing to make any type of edit (insertion, deletion, transition and transversion) mediated by the CRISPR/Cas system(39). However, prime editing is more complex and requires additional refinements compared to base editing. Hence, base editing is considered favorable when possible. BE-FF is currently unable to provide off-targets insights and therefore, such off-targets assessment using complementary tools (e.g. CRISTA(16), Cas-OFFinder(40), CCTop(19) and others) is suggested. For researchers intending to make use of the BE-FF database, in case the mutation of interest is missing, it is suggested to check whether it appears on the full dataset we used. Otherwise, it is recommended to use the web tool to analyze the variation of interest by SNP ID or manual input.

## DATA AVAILABILITY

The **BE-FF** web tool is freely available at: https://www.danioffenlab.com/be-ff

The code is available at: https://github.com/RoyRabinowitz/BE-FF

## SUPPLEMENTARY DATA

- Sup. file1: Dataset of human pathogenic SNPs
- Sup. file2: BE-FF database
- Figure S1

## IMPLEMENTATION

BE-FF is web-based and does not require installation or specific specifications.

## REFERENCES

1. Komor, A.C., Kim, Y.B., Packer, M.S., Zuris, J.A. and Liu, D.R. (2016) Programmable editing of a target base in genomic DNA without double-stranded DNA cleavage. Nature, 533, 420–424.

2. Gaudelli, N.M., Komor, A.C., Rees, H.A., Packer, M.S., Badran, A.H., Bryson, D.I. and Liu, D.R. (2017) Programmable base editing of A•T to G•C in genomic DNA without DNA cleavage. Nature, 551, 464–471.

3. Jinek, M., Chylinski, K., Fonfara, I., Hauer, M., Doudna, J.A. and Charpentier, E. (2012) A programmable dual-RNA-guided DNA endonuclease in adaptive bacterial immunity. Science (80-.)., 337, 816–21.

4. Cong, L., Ran, F.A., Cox, D., Lin, S., Barretto, R., Habib, N., Hsu, P.D., Wu, X., Jiang, W., Marraffini, L.A., et al. (2013) Multiplex genome engineering using CRISPR/Cas systems. Science (80-.)., 339, 819–23.

5. Qi, L.S., Larson, M.H., Gilbert, L.A., Doudna, J.A., Weissman, J.S., Arkin, A.P. and Lim, W.A. (2013) Repurposing CRISPR as an RNA-guided platform for sequence-specific control of gene expression. Cell, 152, 1173–83.

6. Yasui, M., Suenaga, E., Koyama, N., Masutani, C., Hanaoka, F., Gruz, P., Shibutani, S., Nohmi, T., Hayashi, M. and Honma, M. (2008) Miscoding properties of 2’-deoxyinosine, a nitric oxide-derived DNA Adduct, during translesion synthesis catalyzed by human DNA polymerases. J. Mol. Biol., 377, 1015–23.

7. Rees, H.A. and Liu, D.R. (2018) Base editing: precision chemistry on the genome and transcriptome of living cells. Nat. Rev. Genet., 19, 770–788.

8. Ceccaldi, R., Rondinelli, B. and D’Andrea, A.D. (2016) Repair Pathway Choices and Consequences at the Double-Strand Break. Trends Cell Biol., 26, 52–64.

9. Ihry, R.J., Worringer, K.A., Salick, M.R., Frias, E., Ho, D., Theriault, K., Kommineni, S., Chen, J., Sondey, M., Ye, C., et al. (2018) p53 inhibits CRISPR–Cas9 engineering in human pluripotent stem cells. Nat. Med., 24, 939–946.

10. Haapaniemi, E., Botla, S., Persson, J., Schmierer, B. and Taipale, J. (2018) CRISPR–Cas9 genome editing induces a p53-mediated DNA damage response. Nat. Med., 24, 927–930.

11. Kosicki, M., Tomberg, K. and Bradley, A. (2018) Repair of double-strand breaks induced by CRISPR–Cas9 leads to large deletions and complex rearrangements. Nat. Biotechnol., 36, 765–771.

12. Hanlon, K.S., Kleinstiver, B.P., Garcia, S.P., Zaborowski, M.P., Volak, A., Spirig, S.E., Muller, A., Sousa, A.A., Tsai, S.Q., Bengtsson, N.E., et al. (2019) High levels of AAV vector integration into CRISPR-induced DNA breaks. Nat. Commun., 10, 4439.

13. Ran, F.A., Hsu, P.D., Wright, J., Agarwala, V., Scott, D.A. and Zhang, F. (2013) Genome engineering using the CRISPR-Cas9 system. Nat. Protoc., 8, 2281–2308.

14. Yeh, W.-H., Chiang, H., Rees, H.A., Edge, A.S.B. and Liu, D.R. (2018) In vivo base editing of post-mitotic sensory cells. Nat. Commun., 9, 2184.

15. Cong, L., Ran, F.A., Cox, D., Lin, S., Barretto, R., Habib, N., Hsu, P.D., Wu, X., Jiang, W., Marraffini, L.A., et al. (2013) Multiplex Genome Engineering Using CRISPR/Cas Systems. Science (80-.)., 339, 819–823.

16. Abadi, S., Yan, W.X., Amar, D. and Mayrose, I. (2017) A machine learning approach for predicting CRISPR-Cas9 cleavage efficiencies and patterns underlying its mechanism of action. PLoS Comput. Biol., 13, e1005807.

17. Haeussler, M., Schönig, K., Eckert, H., Eschstruth, A., Mianné, J., Renaud, J.-B., Schneider-Maunoury, S., Shkumatava, A., Teboul, L., Kent, J., et al. (2016) Evaluation of off-target and on-target scoring algorithms and integration into the guide RNA selection tool CRISPOR. Genome Biol., 17, 148.

18. Labun, K., Montague, T.G., Krause, M., Torres Cleuren, Y.N., Tjeldnes, H. and Valen, E. (2019) CHOPCHOP v3: expanding the CRISPR web toolbox beyond genome editing. Nucleic Acids Res., 47, W171–W174.

19. Stemmer, M., Thumberger, T., del Sol Keyer, M., Wittbrodt, J. and Mateo, J.L. (2015) CCTop: An Intuitive, Flexible and Reliable CRISPR/Cas9 Target Prediction Tool. PLoS One, 10, e0124633.

20. Hwang, G.H., Park, J., Lim, K., Kim, S., Yu, J., Yu, E., Kim, S.T., Eils, R., Kim, J.S. and Bae, S. (2018) Web-based design and analysis tools for CRISPR base editing. BMC Bioinformatics, 19.

21. Wang, Y., Gao, R., Wu, J., Xiong, Y.C., Wei, J., Zhang, S., Yang, B., Chen, J. and Yang, L. (2019) Comparison of cytosine base editors and development of the BEable-GPS database for targeting pathogenic SNVs. Genome Biol., 20.

22. Rees, H.A., Komor, A.C., Yeh, W.-H., Caetano-Lopes, J., Warman, M., Edge, A.S.B. and Liu, D.R. (2017) Improving the DNA specificity and applicability of base editing through protein engineering and protein delivery. Nat. Commun., 8, 15790.

23. Komor, A.C., Zhao, K.T., Packer, M.S., Gaudelli, N.M., Waterbury, A.L., Koblan, L.W., Kim, Y.B., Badran, A.H. and Liu, D.R. (2017) Improved base excision repair inhibition and bacteriophage Mu Gam protein yields C:G-to-T:A base editors with higher efficiency and product purity. Sci. Adv., 3, eaao4774.

24. Koblan, L.W., Doman, J.L., Wilson, C., Levy, J.M., Tay, T., Newby, G.A., Maianti, J.P., Raguram, A. and Liu, D.R. (2018) Improving cytidine and adenine base editors by expression optimization and ancestral reconstruction. Nat. Biotechnol., 36, 843–846.

25. Kim, Y.B., Komor, A.C., Levy, J.M., Packer, M.S., Zhao, K.T. and Liu, D.R. (2017) Increasing the genome-targeting scope and precision of base editing with engineered Cas9-cytidine deaminase fusions. Nat. Biotechnol., 35, 371–376.

26. Li, X., Wang, Y., Liu, Y., Yang, B., Wang, X., Wei, J., Lu, Z., Zhang, Y., Wu, J., Huang, X., et al. (2018) Base editing with a Cpf1-cytidine deaminase fusion. Nat. Biotechnol., 36, 324–327.

27. Nishida, K., Arazoe, T., Yachie, N., Banno, S., Kakimoto, M., Tabata, M., Mochizuki, M., Miyabe, A., Araki, M., Hara, K.Y., et al. (2016) Targeted nucleotide editing using hybrid prokaryotic and vertebrate adaptive immune systems. Science, 353, aaf8729–aaf8729.

28. Nishimasu, H., Shi, X., Ishiguro, S., Gao, L., Hirano, S., Okazaki, S., Noda, T., Abudayyeh, O.O., Gootenberg, J.S., Mori, H., et al. (2018) Engineered CRISPR-Cas9 nuclease with expanded targeting space. Science, 361, 1259–1262.

29. Hu, J.H., Miller, S.M., Geurts, M.H., Tang, W., Chen, L., Sun, N., Zeina, C.M., Gao, X., Rees, H.A., Lin, Z., et al. (2018) Evolved Cas9 variants with broad PAM compatibility and high DNA specificity. Nature, 556, 57–63.

30. Gehrke, J.M., Cervantes, O., Clement, M.K., Wu, Y., Zeng, J., Bauer, D.E., Pinello, L. and Joung, J.K. (2018) An APOBEC3A-Cas9 base editor with minimized bystander and off-target activities. Nat. Biotechnol., 36, 977–982.

31. Jiang, W., Feng, S., Huang, S., Yu, W., Li, G., Yang, G., Liu, Y., Zhang, Y., Zhang, L., Hou, Y., et al. (2018) BE-PLUS: a new base editing tool with broadened editing window and enhanced fidelity. Cell Res., 28, 855–861.

32. Oakes, B.L., Fellmann, C., Rishi, H., Taylor, K.L., Ren, S.M., Nadler, D.C., Yokoo, R., Arkin, A.P., Doudna, J.A. and Savage, D.F. (2019) CRISPR-Cas9 Circular Permutants as Programmable Scaffolds for Genome Modification. Cell, 176, 254-267.e16.

33. Huang, T.P., Zhao, K.T., Miller, S.M., Gaudelli, N.M., Oakes, B.L., Fellmann, C., Savage, D.F. and Liu, D.R. (2019) Circularly permuted and PAM-modified Cas9 variants broaden the targeting scope of base editors. Nat. Biotechnol., 37, 626–631.

34. Thuronyi, B.W., Koblan, L.W., Levy, J.M., Yeh, W.-H., Zheng, C., Newby, G.A., Wilson, C., Bhaumik, M., Shubina-Oleinik, O., Holt, J.R., et al. (2019) Continuous evolution of base editors with expanded target compatibility and improved activity. Nat. Biotechnol., 37, 1070–1079.

35. Ryu, S.-M., Koo, T., Kim, K., Lim, K., Baek, G., Kim, S.-T., Kim, H.S., Kim, D.-E., Lee, H., Chung, E., et al. (2018) Adenine base editing in mouse embryos and an adult mouse model of Duchenne muscular dystrophy. Nat. Biotechnol., 36, 536–539.

36. Hua, K., Tao, X. and Zhu, J.-K. (2019) Expanding the base editing scope in rice by using Cas9 variants. Plant Biotechnol. J., 17, 499–504.

37. Yang, L., Zhang, X., Wang, L., Yin, S., Zhu, B., Xie, L., Duan, Q., Hu, H., Zheng, R., Wei, Y., et al. (2018) Increasing targeting scope of adenosine base editors in mouse and rat embryos through fusion of TadA deaminase with Cas9 variants. Protein Cell, 9, 814–819.

38. Sherry, S.T., Ward, M. and Sirotkin, K. (1999) dbSNP - database for single nucleotide polymorphisms and other classes of minor genetic variation. Genome Res., 9, 677–679.

39. Anzalone, A. V, Randolph, P.B., Davis, J.R., Sousa, A.A., Koblan, L.W., Levy, J.M., Chen, P.J., Wilson, C., Newby, G.A., Raguram, A., et al. (2019) Search-and-replace genome editing without double-strand breaks or donor DNA. Nature, 10.1038/s41586-019-1711-4.

40. Bae, S., Park, J. and Kim, J.-S. (2014) Cas-OFFinder: a fast and versatile algorithm that searches for potential off-target sites of Cas9 RNA-guided endonucleases. Bioinformatics, 30, 1473–1475.

